# Orchestration of Chlorophyll and Carotenoid Biosynthesis by ORANGE Family Proteins in Plant

**DOI:** 10.1101/2022.02.08.479616

**Authors:** Tianhu Sun, Peng Wang, Shan Lu, Hui Yuan, Yong Yang, Tara Fish, Theodore Thannhauser, Jiping Liu, Michael Mazourek, Bernhard Grimm, Li Li

## Abstract

Chlorophyll and carotenoid are essential photosynthetic pigments. Plants must spatiotemporally coordinate the needs of chlorophyll and carotenoid for optimal photosynthesis and plant fitness in response to diverse environmental and developmental cues. However, how these two biosynthesis pathways are orchestrated remains largely unknown. Here, we report that the highly conserved ORANGE (OR) family proteins are the common regulators of both pathways via posttranslationally regulating the first committed enzyme in each pathway. We demonstrate that OR family proteins physically interact with magnesium chelatase subunit I (CHLI) in addition to phytoene synthase (PSY) and concurrently regulate CHLI and PSY protein stability and activity. We show that loss of *OR* genes hinders both chlorophyll and carotenoid biosynthesis, limits light-harvesting complex assembly, and impairs thylakoid grana stacking in chloroplast. *OR* overexpression safeguards photosynthetic pigment biosynthesis and enhances thermotolerance in both Arabidopsis and tomato plants. Our findings establish a conserved mechanism of green plant to coordinate chlorophyll and carotenoid biosynthesis and provide a potential genetic target to generate climate-resilient crops.

## Introduction

Chlorophyll and carotenoid are essential pigments for photosynthesis, making them indispensable to plant and life on the earth. In chloroplast, chlorophyll plays a vital role in converting light energy to chemical energy (Tanaka and Tanaka, 2007; Wang and Grimm, 2021). Carotenoid is accessory light-harvesting pigment and protects the photosynthetic apparatus against photo-oxidative damage by quenching chlorophyll-excited states, scavenging reactive oxygen species, and dissipating excess energy (Niyogi and Truong, 2013; Murchie and Ruban, 2020). Both chlorophyll and carotenoid are obligatory for the structural assembly and stability of photosynthetic apparatus. Plant spatiotemporally coordinates the needs for chlorophyll and carotenoid in efficient photosynthesis and photoprotection by controlling of their biosynthesis pathways. The dysfunction of either pathway impacts the other and severely hinders autotrophic growth (Härtel and Grimm, 1998; Qin et al., 2007). As such, plant has evolved multifaceted mechanisms to orchestrate chlorophyll and carotenoid biosynthesis, which is crucial for optimal photosynthesis, chloroplast development, plant fitness, and crop yield in response to diverse developmental and environmental cues.

The biosynthesis of chlorophyll uses 5-aminolevulinic acid (ALA) to form precursor protoporphyrin IX (ProtoIX) (Figure 1A). ProtoIX is directed to the Mg branch by Mg-chelatase (MgCh), the first committed enzyme of the specific chlorophyll biosynthesis pathway (Tanaka and Tanaka, 2007; Wang and Grimm, 2021). MgCh consists of three subunits, two AAA+ ATPases CHLI and CHLD, and the catalytic subunit CHLH (also called GENOMES UNCOUPLED 5, GUN5). MgCh catalyzes the two-step reaction of catalytic activation and Mg^2+^ insertion into ProtoIX to form Mg protoporphyrin IX (MgProtoIX), and is activated by GUN4 (Jensen et al., 1999; Larkin et al., 2003; Peter and Grimm, 2009; Richter et al., 2016). The subsequent reactions convert MgProtoIX into MgProtoIX monomethylester (MgPMME), and then in continuation into divinyl protochlorophyllide (Pchlide). A light-dependent reaction converts Pchlide into chlorophyllide (Chlide), which is the substrate for the formation of chlorophyll *a* and *b* (Figure 1A). The biosynthesis of carotenoid, a group of tetraterpenoids, starts with glyceraldehyde 3-phosphate and pyruvate to generate the direct precursor geranylgeranyl pyrophosphate (GGPP). Phytoene synthase (PSY) catalyzes the condensation of two GGPPs into phytoene, which represents the first committed and a major rate-limiting step for carotenoid biosynthesis (Cazzonelli and Pogson, 2010; Sun et al., 2018; Sun and Li, 2020).

**Figure 1.**
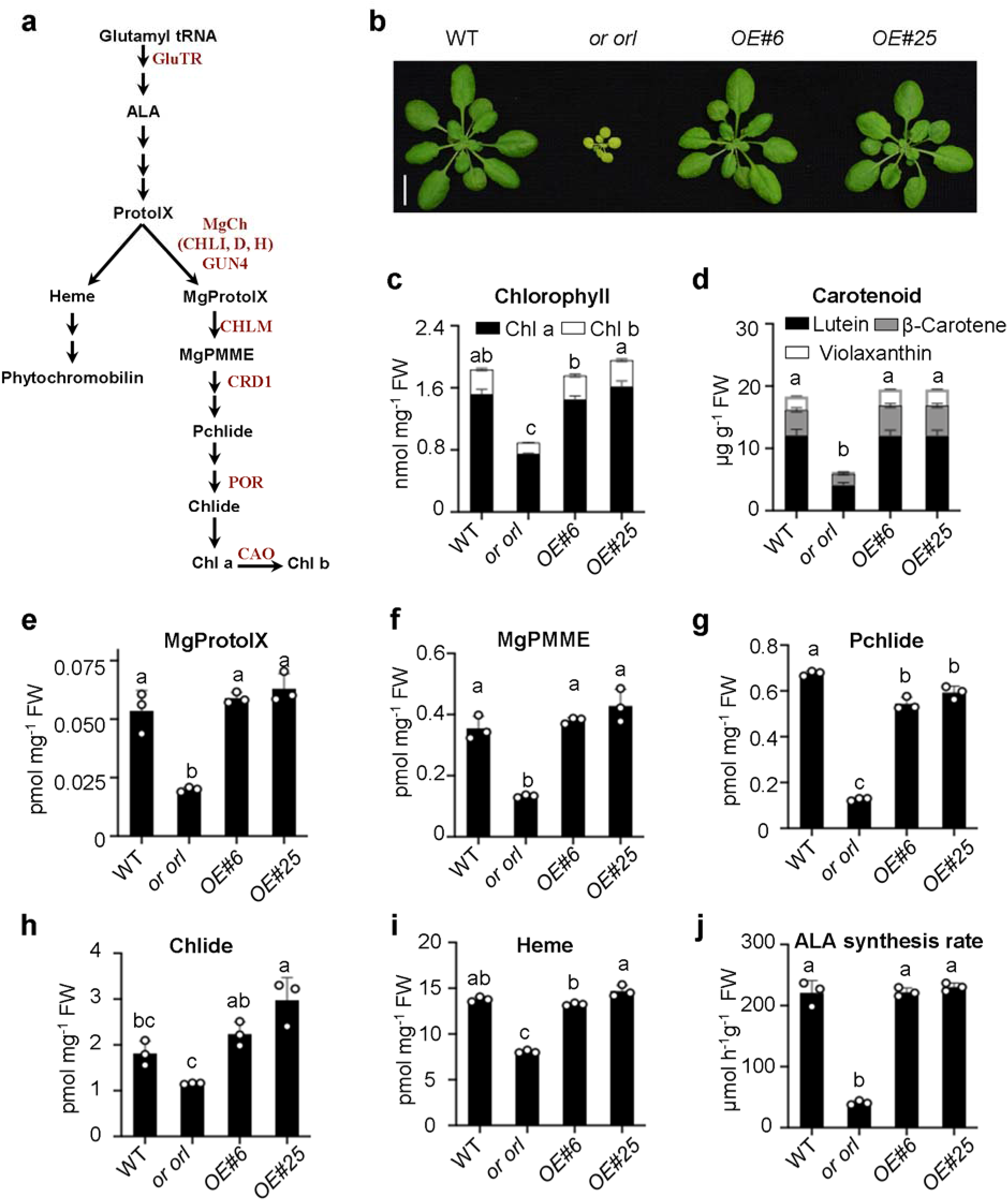
OR proteins affect chlorophyll and carotenoid biosynthesis. **A**, Scheme of chlorophyll biosynthesis pathway in plant. The genes/enzymes investigated in this study are shown in the pathway. GluTR, glutamyl-tRNA reductase; ALA, 5-aminolevulinic acid; ProtoIX, protoporphyrin IX; MgCh, magnesium chelatase; GUN4, GENOMES UNCOUPLED 4; MgProtoIX, magnesium protoporphyrin IX; CHLM, MgProtoIX methyltransferase; MgPMME, MgProtoIX monomethylester; CRD1, MgPMME cyclase 1; Pchlide, protochlorophyllide; POR, Pchlide oxidoreductase; Chlide, chlorophyllide; Chl, chlorophyll; CAO, chlorophyll a oxygenase. **B,** A representative image of 4-week-old Col-0 wild-type (WT), *or orl* double mutant, and *OR* overexpressors (OE#6 and OE#25) grown under 10 h light and 14 h dark, 100 μmol photons per m2 per s. Scale bar, 1 cm. **C,** Chlorophyll level. **D,** Carotenoid level. **E-H,** Levels of chlorophyll biosynthetic intermediates MgProtoIX (**E**), MgPMME (**F**), Pchlide (**G**), and Chlide (**H**), as well as of heme (**I**) and ALA (**J**) at 4-week-old stage of WT, *or orl*, and *OE* lines. **C-J,** Data represent means ± SD, n=3. Letters above histograms indicate significant differences as determined by Tukey’s multiple comparisons test.

Because of the central roles of MgCh and PSY in directing metabolic fluxes into chlorophyll and carotenoid biosynthesis, respectively, both MgCh and PSY are multifacetedly regulated (Tanaka and Tanaka, 2007; Mochizuki et al., 2010; Brzezowski et al., 2015; Nisar et al., 2015; Llorente et al., 2017; Sun and Li, 2020). While transcriptional regulation is important, a range of posttranslational mechanisms allows rapid regulation of MgCh or PSY to affect chlorophyll or carotenoid biosynthesis (Larkin et al., 2003; Zhou et al., 2015; Álvarez et al., 2016; Welsch et al., 2018; Wang et al., 2020; Zhang et al., 2020).

It is well known that the biosynthesis of chlorophyll and carotenoid is under tight and sophisticated regulation in chloroplast to ensure optimal photosynthesis and adaptation (Cazzonelli and Pogson, 2010; Ruiz-Sola and Rodríguez-Concepción, 2012; Kobayashi and Masuda, 2016; Sun and Li, 2020; Wang and Grimm, 2021). However, how these two biosynthesis pathways are coordinately regulated remains poorly understood. The common regulators that orchestrate chlorophyll and carotenoid biosynthesis at the posttranslational level, allowing rapid regulation of these pathways, are unknown.

Regulation of the biosynthesis pathway frequently occurs at the first committed step of the pathway. DnaJE1 type chaperone ORANGE (OR) family proteins OR and OR-Like (ORL) are the master regulators of PSY to control carotenoid biosynthesis in chloroplast (Zhou et al., 2015). In this study, we discover that the OR family proteins posttranslationally regulate CHLI, an essential subunit of MgCh, to mediate chlorophyll biosynthesis in addition to regulating PSY for carotenoid biosynthesis. OR family proteins serve as common conserved regulators of chlorophyll and carotenoid biosynthesis, and affect chloroplast development and thermotolerance in plant.

## Results

### OR family proteins directly interact with Mg-chelatase subunit I to affect chlorophyll biosynthesis

In our previous study, we observed that the *or orl* double knock-out mutant has dramatically reduced photosynthetic pigments compared to the wild type *Arabidopsis thaliana* (WT) (Zhou et al., 2015). The *or orl* mutant showed retarded growth and pale-green leaves (Figure 1B) with reduced chlorophyll level (Figure 1C) and decreased carotenoid content (Figure 1D). To investigate whether the OR family proteins play a role in chlorophyll biosynthesis, the levels of the chlorophyll metabolic intermediates in WT, *or orl*, and *OR* overexpression (*OE*) lines were measured. The steady state levels of Mg-porphyrins (including MgProtoIX and MgPMME) decreased significantly in *or orl* in comparison with WT (Figure 1, E and F). Similarly, the content of downstream metabolites Pchlide and Chlide were also reduced in the mutant (Figure 1, G and H). Interestingly, the heme branch of tetrapyrrole was also disturbed (Figure 1I), probably owing to the drastically reduced ALA synthesis rate in the *or orl* mutant (Figure 1J). The *OE* lines did not significantly accumulate more of these metabolites, with the exception of the Chlide content (Figure 1).

The accumulation of intermediates caused by impaired downstream reactions in the chlorophyll biosynthesis pathway can be visualized *in vivo* by the specific emission spectrum (Ankele et al., 2007). To see whether a specific step in the chlorophyll biosynthesis pathway is affected by the OR proteins, an ALA feeding experiment was carried out in *or orl* along with several mutants (i.e. *chli, gun5, crd1*, and *cao*) in the chlorophyll biosynthesis pathway (Figure 1A). All those mutant lines showed retarded growth with a pale green leaf phenotype (Figure 2A). Consistent with a previous report (Ankele et al., 2007), the ALA-fed *chli* and *gun5* accumulated ProtoIX because of the impaired MgCh activity. Remarkably, *or orl* also accumulated ProtoIX to a higher extent than WT (Figure 2B), indicating that the OR family proteins affect MgCh activity. In contrast, *crd1* and *cao* showed similar ProtoIX level as WT control.

**Figure 2.**
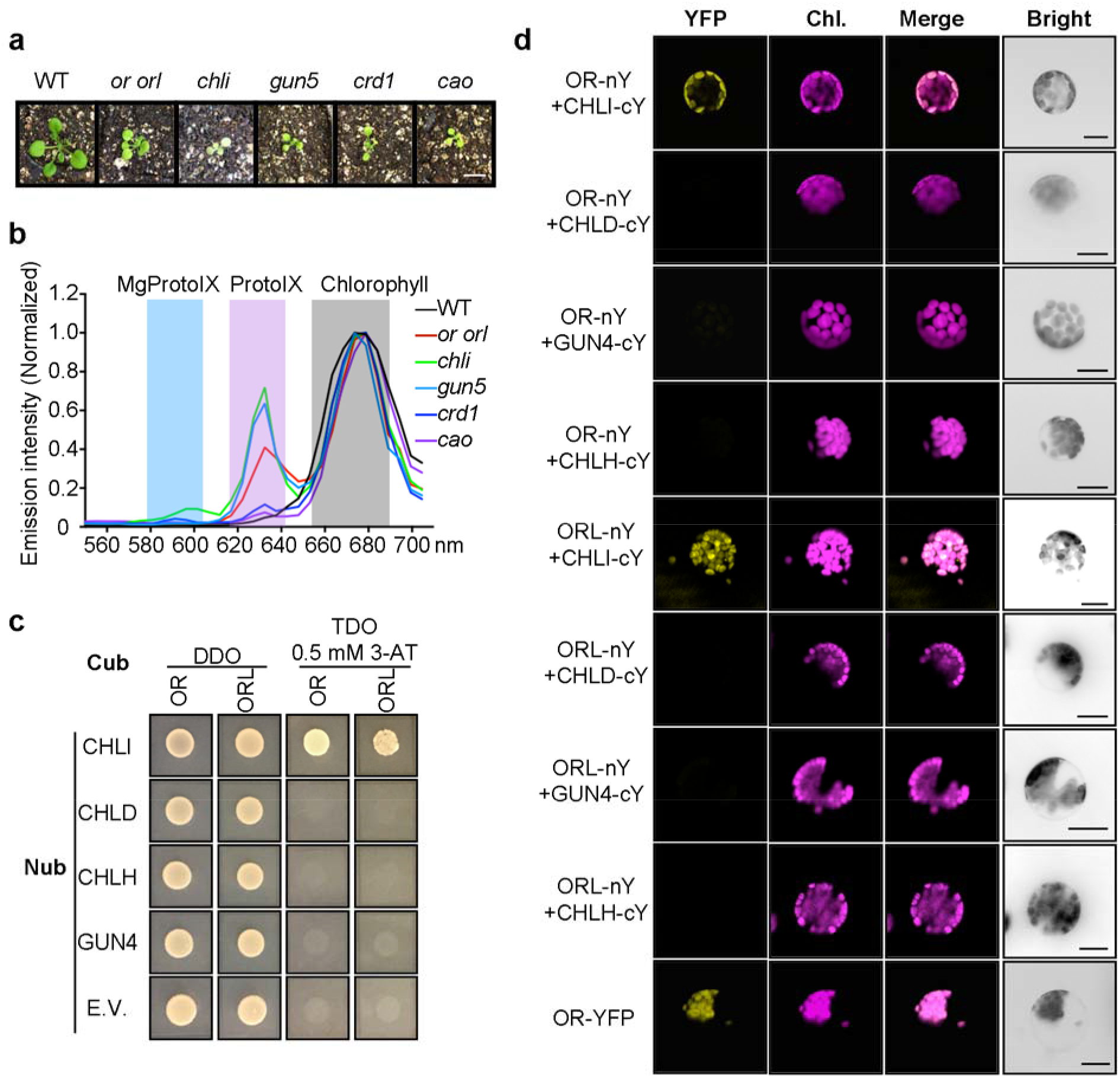
OR family proteins directly interact with CHLI. **A**, Representative images of 14-day-old WT, *or orl, chli, gun5, crd1*, and *cao* mutant plants. Scale bar, 0.5 cm. **B,** Emission spectra scan of 6-day-old seedlings fed with ALA in dark using confocal laser scanning microscopy. The emission peaks at 585 to 615 nm, 627 to 640 nm, and 660 to 710 nm are highlighted for the specific emission of Mg-porphyrins (including MgProtoIX and MgPMME), ProtoIX, and chlorophyll. The emission intensity was normalized to the maximum value for each measurement, and calculated by averaging measurements of five positions overlapping chloroplasts. **C,** Y2H analysis of interactions between OR family proteins and MgCh subunits (CHLI, CHLD, CHLH) and GUN4. The transformed yeast strains were grown on medium lacking Leu and Trp (DDO) or His, Leu, and Trp (TDO) with 0.5 mM 3-amino-1,2,4-triazole (3-AT). The Nub empty vector (E.V.) was used as negative control. **D,** BiFC assay of interactions between OR proteins and MgCh-related proteins in Arabidopsis protoplasts. The expression of OR-YFP served as the positive control. Chl, chlorophyll autofluorescence; Scale bars, 10 μm.

OR has been shown to regulate carotenoid biosynthesis by physically interacting with PSY in Arabidopsis (Zhou et al., 2015). Thus, we investigated the possible interactions between the OR family proteins and chlorophyll biosynthetic enzymes (Supplemental Figure S1). As ProtoIX accumulation in *or orl* pointed to impaired Mg chelation, we focused on the interactions between OR proteins and the three subunits of MgCh and GUN4. Both OR and ORL physically interacted with CHLI of MgCh in the Y2H analysis (Figure 2C, Supplemental Figure S1). Further, the bimolecular fluorescence complementation (BiFC) assay in the protoplasts evidenced a direct interaction of OR and ORL with CHLI in chloroplast (Figure 2D). No interactions of OR or ORL with CHLD, CHLH, GUN4, CHLM, CRD1, or PORB were confirmed by both assays (Figure 2C and 2D, Supplemental Figure S1 and S2). These findings suggest that in addition to mediating carotenoid biosynthesis by physically interacting with PSY (Zhou et al., 2015), OR family proteins also directly regulate chlorophyll biosynthesis through interaction with CHLI, an essential component of the first committed enzyme in the chlorophyll biosynthesis pathway.

The sub-organellar localization of CHLI and PSY is ambiguous since these two proteins are reported in both soluble and membrane fractions of chloroplast (Joyard et al., 2009; Shumskaya et al., 2012). However, the active form of both MgCh and PSY requires membrane association in chloroplast (Welsch et al., 2000; Adhikari et al., 2009; Welsch et al., 2018). Immunoblot analysis of isolated chloroplast envelope, stroma, and thylakoid fractions showed that OR protein was mainly located in the thylakoid membrane fraction (Supplemental Figure S3). Similarly, CHLI and PSY protein were also dominantly presented in the same location (Supplemental Figure S3). Therefore, the intrinsic membrane association of OR could enable the coordinated regulation of CHLI and PSY, which predominantly function in their membrane-associated forms.

### Specific effects of OR proteins on light-harvesting complex assembly and thylakoid membrane stacking

Both chlorophyll and carotenoid are the essential constituents of photosynthetic complexes and are required for chloroplast development. To investigate the effects of *OR* proteins on the photosynthetic complex formation, we performed blue native polyacrylamide gel electrophoresis (BN-PAGE). In comparison with WT control, the *or orl* mutant had less light-harvesting complex II (LHCII) assembly along with less PSI+PSII dimer and super complex formation (Figure 3A).

**Figure 3.**
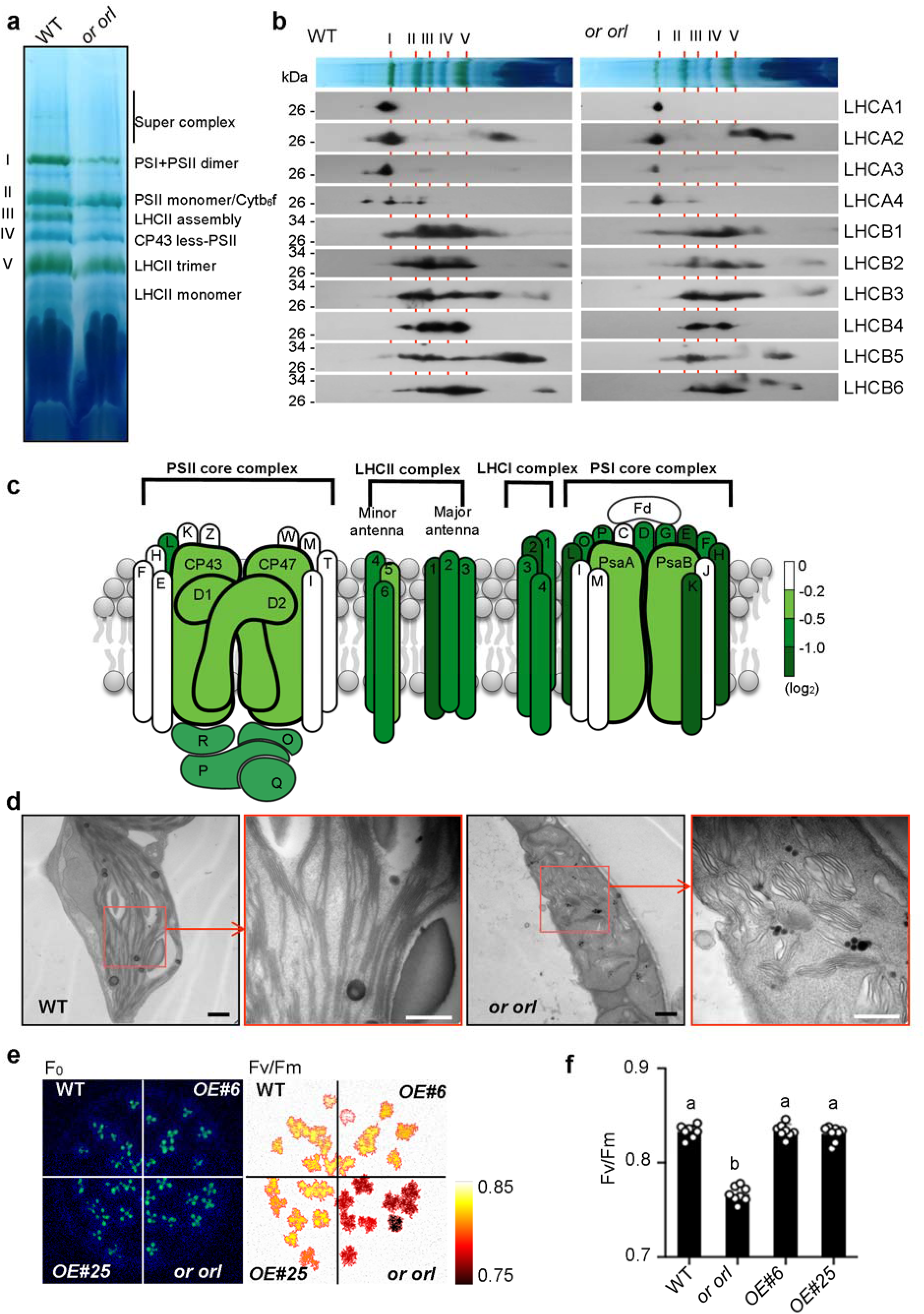
OR proteins are necessary for light harvesting complex assembly and thylakoid membrane stacking. **A**, A representative BN-PAGE gel of thylakoid membrane pigment-protein complexes from WT and *or orl* mutant. Equal amounts of thylakoid membranes (8 μg of chlorophyll) from WT and *or orl* plants were solubilized in 1% (w/v) Triton X-100. The major complexes (I, II, III, IV, and V) are determined according to previous reports (Wang and Grimm, 2016). **B,** Two-dimensional BN/SDS-PAGE and immunoblot analysis of LHC proteins. Individual lanes from the BN-PAGE gel in (**A**) were subjected to SDS-PAGE and immunoblot with corresponding antibodies in the second dimension. **C,** Proteomic profiling of photosystem proteins in *or orl* mutant in comparison to WT. Log ratios are used for visualization of fold changes (Supplemental Table S2). **D,** Electron micrographs of leaf chloroplasts of 3-week-old plants of WT and the *or orl* mutant. Scale bars, 500 nm. **E-F,** Measurement of chlorophyll fluorescence parameters of WT, *or orl*, and *OE* line seedlings germinated on agar plates. False-color images represented F0 and Fv/Fm (**E**), Fv/Fm was also presented in (**F**) as means ± SE, n=8, Letters above histograms indicate significant differences as determined by Tukey’s multiple comparisons test.

Light-harvesting complexes comprise LHC proteins and the assembled photosynthetic pigments. By combining of two-dimensional BN/SDS-PAGE with immunoblot analysis, we found that the reduction of photosynthetic pigments in *or orl* led to reduced levels of LHC proteins, particularly LHCB1 and LHCB2, in the complexes (Figure 3B, Supplemental Figure S4). Moreover, the total steady state levels of several LHC proteins were reduced in *or orl* in comparison with WT (Supplemental Figure S5). Additional proteomics analysis unveiled that the LHC antenna assembly and some photosystem core proteins had reduced protein levels in *or orl* (Figure 3C, Table S2). These findings were consistent with the high correlation between impaired synthesis of pigments and low LHC protein content (Dall’Osto et al., 2015; Wang and Grimm, 2021).

The LHCII along with the LHCII-PSII supercomplexes is important for the formation of grana stacks in chloroplast (Albanese et al., 2020). To see how the defect of light-harvesting complex assembly in *or orl* affects grana formation and chloroplast development, chloroplast ultrastructure was examined by transmission electron microscopy. Chloroplast of WT had the typical multiple stacked layers of granular thylakoids interconnected by stroma lamellae (Figure 3D). However, chloroplast of the *or orl* mutant had abolished granular stacks, disrupted stroma lamellae, and swollen membrane structure, showing distinct impairment and abnormal ultrastructural organization of thylakoid membranes (Figure 3D). Notably, this defect of thylakoid membrane ultrastructure resembled that in the *chli1* mutant (Apchelimov et al., 2007; Myouga et al., 2013).

Chlorophyll fluorescence parameters Fv/Fm (variable chlorophyll fluorescence/maximal fluorescence of dark-adapted leaves) represent the maximum photochemical efficiency of PSII (Murchie and Lawson, 2013). The effect of photosynthetic pigment alteration in *or orl* and *OE* lines on photosynthesis were also assessed in comparison with WT. While *OE* plants had no difference in the Fv/Fm ratio compared to WT, the *or orl* mutant showed a lower Fv/Fm ratio (Figure 3, E and F). Taken together, these findings reveal the impacts of OR-regulated photosynthetic pigment synthesis on LHCII assembly, thylakoid membrane development in chloroplast, and photosynthesis efficiency.

### OR proteins concurrently regulate CHLI and PSY protein stability

OR family proteins were found to physically interact with CHLI (Figure 2). To investigate how such interaction influences chlorophyll biosynthesis, expression of *CHLI* and several chlorophyll biosynthesis related genes (i.e. *GluTR, CHLD, CHLH*, and *GUN4*) along with *PSY* were first examined in WT, *or orl*, and *OE* lines. No significant changes in the transcript levels of these genes were observed (Figure 4A). Therefore, the defect in photosynthetic pigment synthesis in *or orl* is unlikely caused by reduced expression of these genes.

**Figure 4.**
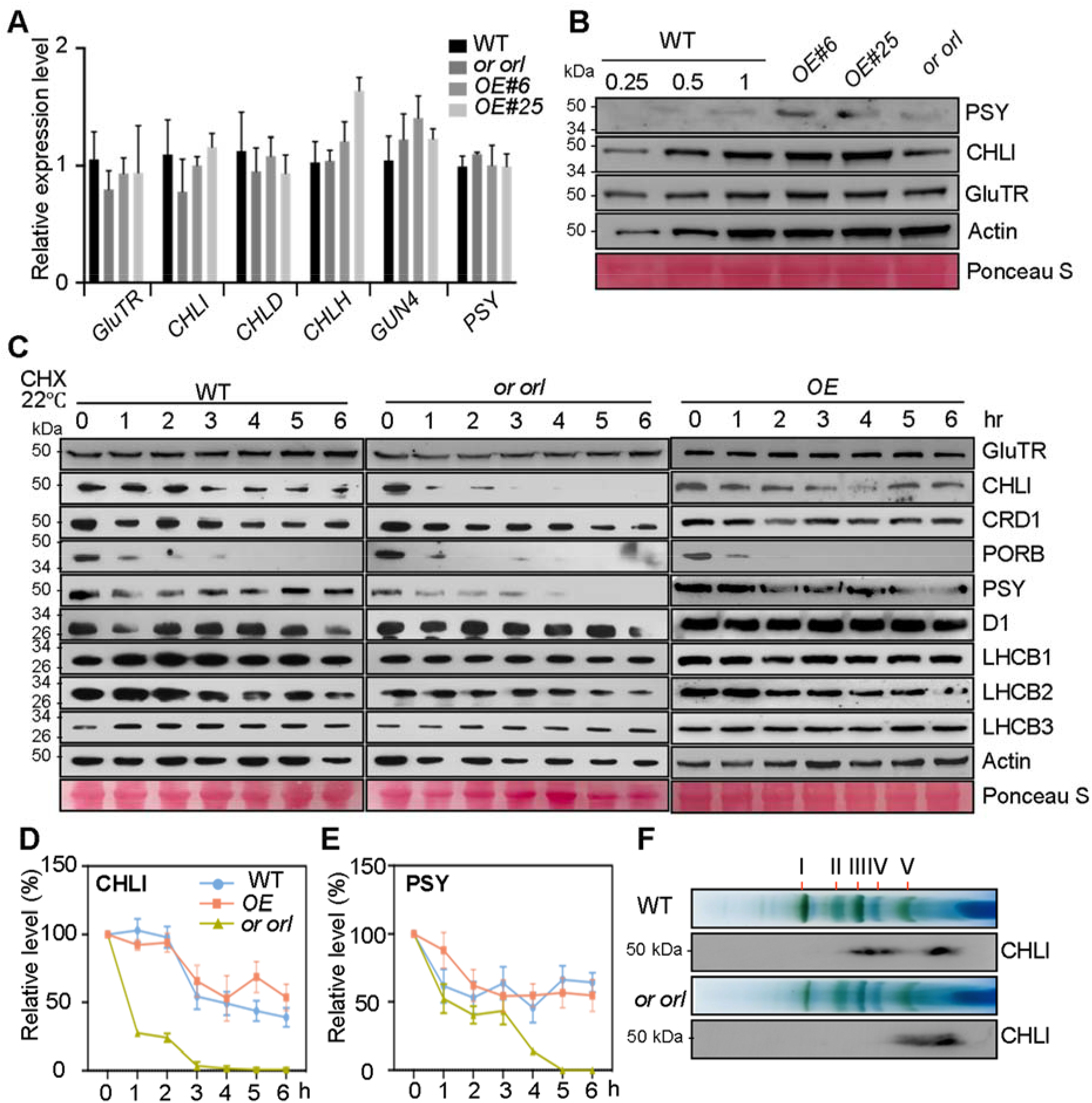
OR proteins concurrently regulate the steady state protein levels of CHLI and PSY. **A,** Relative expression of genes in 4-week-old plants quantified by real-time PCR. Data are means ± SE, n=3. **B,** Steady state protein levels in 4-week-old plants by immunoblot analysis with the indicated antibodies. Proteins from WT were loaded with a series of dilutions for comparison. **C,** Immunoblot analysis of various protein turnover after 100 μmol cycloheximide (CHX) treatment at 22 °C to inhibit new protein synthesis. Relative CHLI (**D**) and PSY (**E**) protein levels were normalized and expressed as the percentage relative to the levels at 0 hr as quantified from immunoblot images. Data represent the means ± SD from three biological replicates. **F,** Two-dimensional BN/SDS-PAGE and immunoblot analysis of CHLI protein from WT and *or orl* thylakoid membranes. The major bands on BN-PAGE gel were labeled same as in Figure 3a.

Further immunoblot analysis revealed that the steady state level of CHLI appeared to be slightly up-regulated in *OE* lines, but down-regulated in *or orl* (Figure 4B). The PSY level was increased in the *OE* lines but reduced in *or orl*, consistent with our previous observation (Zhou et al., 2015). In addition, we examined the level of glutamyl-tRNA reductase (GluTR), the rate-limiting enzyme of ALA synthesis. The GluTR protein level was similar in all lines examined (Figure 4B).

OR is a chaperone protein with holdase activity (Park et al., 2016; D’Andrea et al., 2018). Chaperones bind to client proteins to prevent aggregation and assist in refolding, thereby enhancing their stability. To see whether CHLI is a client protein of OR, a protein sedimentation assay was performed. A significant proportion (31% and 27%, respectively) of both aggregated His-CHLI and GST-CHLI were refolded and presented in a soluble fraction when mixed with the GST-fused N-terminal domain of OR (OR-N), documenting CHLI as a client protein of OR (Supplemental Figure S6).

To investigate whether OR regulates CHLI protein stability, the leaves of WT, *or orl*, and *OE* lines were treated with cycloheximide (CHX), which inhibits *de novo* protein translation. While CHLI was detected after a 6-h-treatment in WT control, it was undetectable after a 3-h-treatment in *or orl*, pointing to a faster turnover of the CHLI protein (Figure 4, C and D). In contrast, other selected chlorophyll biosynthetic enzymes, such as GluTR, CRD1, and PORB, and photosystem components, such as D1 and LHCB1-3, exhibited no difference in stability in *or orl* in comparison with WT control (Figure 4C). A fast turnover of PSY was also found in *or orl* (Figure 4, C and E). These findings demonstrate an essential role of OR family proteins in concurrently regulating both CHLI and PSY stability for chlorophyll and carotenoid biosynthesis, respectively.

MgCh functions after assembly of the subunits CHLI, CHLD, and CHLH (GUN5), and is activated by GUN4 (Larkin et al., 2003; Peter and Grimm, 2009; Richter et al., 2016). We also examined and compared the integrity of the CHLI-containing polymeric complex assembling in *or orl* and WT. In WT chloroplast, CHLI was detected in a large complex as well as in a free form via immunoblot analysis of BN/SDS-PAGE (Figure 4F). However, the large assembled complex containing CHLI in *or orl* disappeared, implying a defect in forming the CHLI-containing protein complex. It is likely that the lack of OR family proteins perturbs the CHLI contribution to the active form of MgCh.

### OR mediates MgCh and PSY enzymatic activity to safeguard photosynthetic pigment synthesis under heat stress

Among the chlorophyll biosynthesis enzymes, CHLI has been documented as one of the proteins most affected by heat stress (Rocco et al., 2013; Echevarría-Zomeño et al., 2016). Therefore, the protein stability of CHLI along with PSY and LHCB1-3 was further examined in plants treated at 37 °C. As expected, faster turnovers of CHLI and PSY proteins were observed in WT after the heat treatment compared to the normal growth temperature at 22 °C (Figure 4C and Figure 5A). Under the heat stress at 37 °C, CHLI and PSY degraded after 3 hrs of treatment in WT, whereas these proteins were barely detectable in *or orl* after 1.5 hrs (Figure 5A). In contrast, CHLI and PSY proteins were persistently observed during the heat treatment in *OE* lines (Figure 5, A, B, and C). These results document that OR proteins simultaneously promote CHLI and PSY stability at elevated temperature.

**Figure 5.**
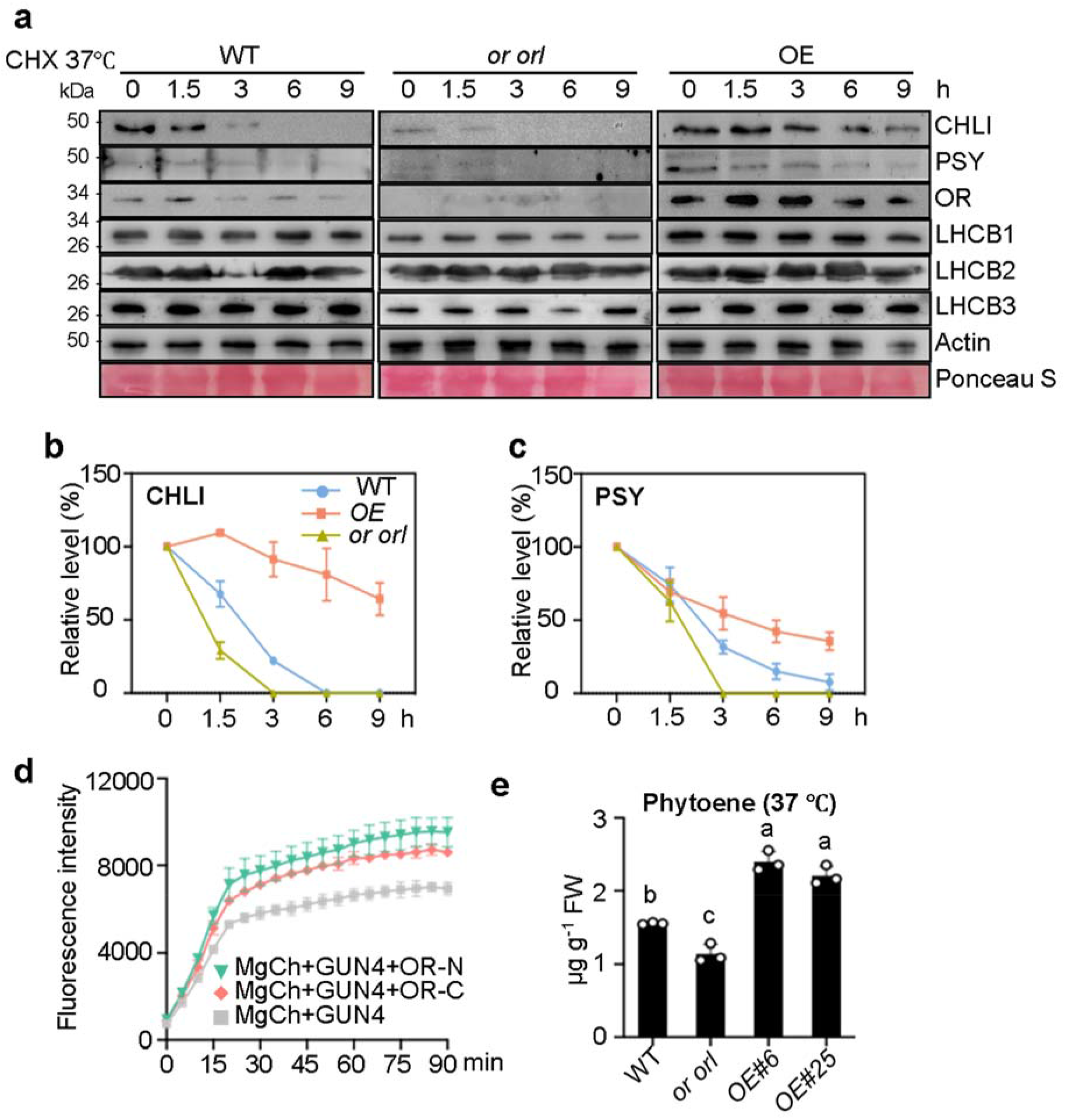
OR proteins maintain PSY and CHLI protein stability and enzymatic activity under heat stress. **A,** Immunoblot analysis of various protein turnover after 100 μmol CHX treatment at 37 °C; Relative CHLI (**B**) and PSY (**C**) protein levels were normalized and expressed as the percentage relative to the levels at 0 hr as quantified from immunoblot images. Data represent the means ± SD from three biological replicates. **D,** *In vitro* MgCh assay. Production of MgProtoIX in the assay was quantified by the fluorescence intensity. Recombinant CHLI, CHLD, CHLH, GUN4, OR-N, and OR-C proteins were used in the assay as indicated. Data represent means ± SD. **E,** Measurement of PSY activity at 37 °C by quantification of phytoene level following norflurazon treatment. Data represent means ± SD, n=3. Letters above histograms indicate significant differences as determined by Tukey’s multiple comparisons test.

We next examined MgCh activity by quantifying the production of MgProtoIX with fluorescence (Ji et al., 2021) using the recombinant MgCh subunits (CHLI, CHLD, and CHLH) and GUN4 with or without OR protein (Supplemental Figure S7). At room temperature, the addition of OR was not observed to affect the MgCh activity (Supplemental Figure S8). However, after a 5 min heat shock treatment of the reaction mixture at 42 °C, OR promoted significantly higher MgCh activity in comparison to the Mg chelation in the absence of OR (Figure 5D), showing a role of OR in regulating MgCh activity.

We also examined PSY activity by measuring phytoene levels following treatment of plant with norflurazon, an inhibitor of phytoene desaturase, to block further metabolism of phytoene. The accumulation of phytoene level is known to directly reflect PSY activity and the method is commonly used to measure PSY activity *in vivo* (Rodríguez-Villalón et al., 2009; Welsch et al., 2018). Upon norflurazon treatment of the plants grown under a heat stress condition at 37 °C, the *OE* lines contained significantly higher phytoene level than WT control (Figure 5E). Taken together, these results showed that OR safeguards both MgCh and PSY activity for photosynthetic pigment synthesis under heat stress.

### Overexpression of *OR* enhances plant thermotolerance

Since OR safeguarded photosynthetic pigment synthesis at elevated temperature (Figure 5), the thermotolerance of several *OE* lines (Supplemental Figure S9) and the *or orl* mutant was assessed. Under the normal growth temperature at 22 °C, the *or orl* plant showed reduced biomass, whereas the *OE* lines were comparable to WT control (Figure 6, A and B). Following a 48 hour treatment at 37 °C and subsequent recovery for 48 hours, the WT plants lost 38.6% of fresh weight while the *or orl* plants displayed a 62% reduction in fresh weight with severe leaf chlorosis in comparison with the lines growing at 22 °C. In contrast, the *OE* plants were less affected (Figure 6A) and lost only 17.3% to 20.6% of fresh weight, resulting in significantly higher biomass compared to WT (Figure 6B). Moreover, compared to over a 50% reduction in pigment content in WT, the *OE* plants had less impaired chlorophyll and carotenoid accumulation with significantly higher pigment level than WT control under heat stress (Figure 6C). These results indicate the importance of OR family proteins in protecting plants from heat stress by safeguarding photosynthetic pigment synthesis.

**Figure 6.**
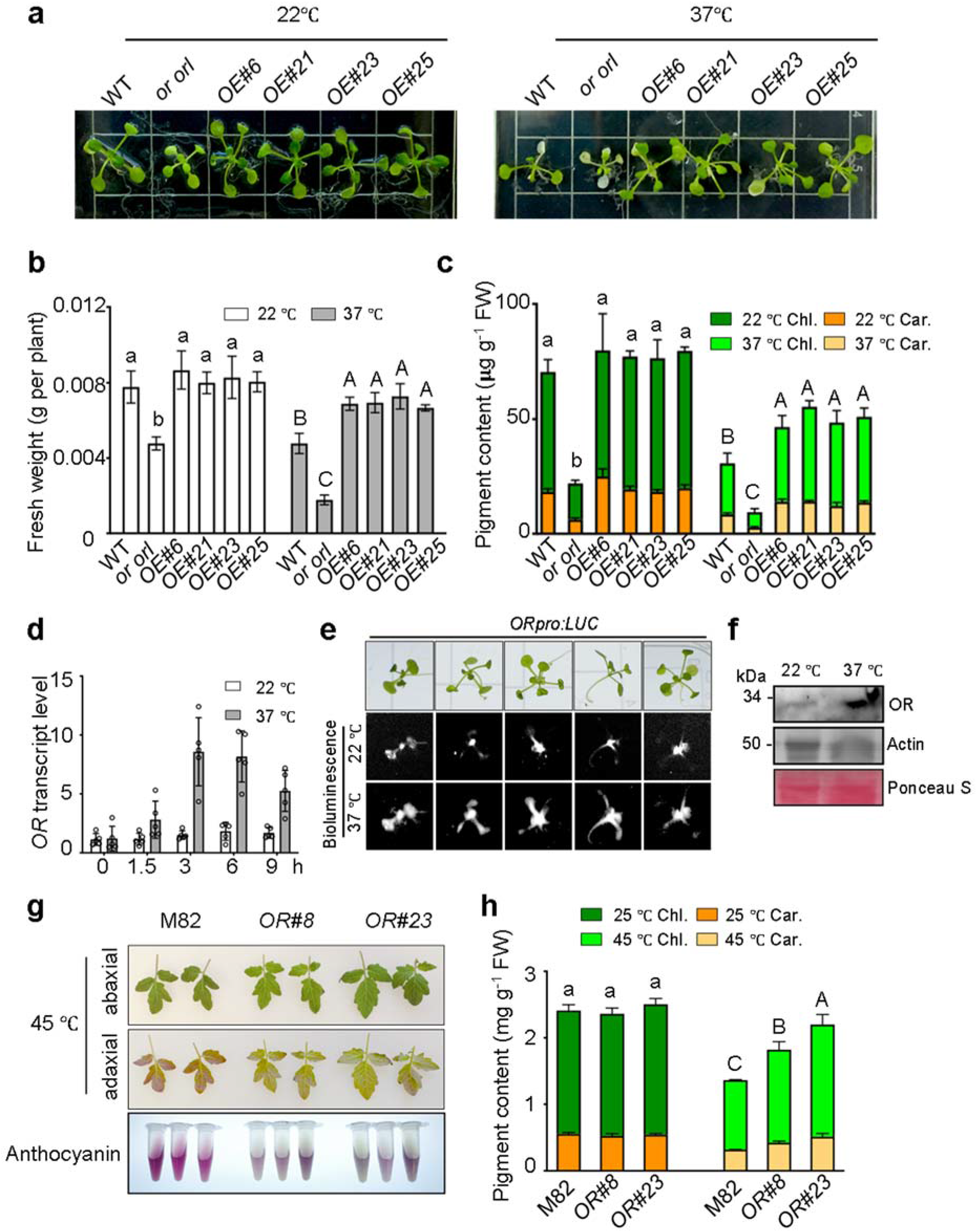
OR proteins affect plant thermotolerance. **A**, Representative phenotype of two-week-old WT, *or orl* and *OE* lines grown on plates at 22 °C as control and following heat treatment at 37 °C for 48 hours and then recovery for 48 hours. **B,** Fresh weight of plants grown at 22 °C and 37 °C. **C,** Chlorophyll (Chl) and carotenoid (Car) pigment content of plants grown at 22 °C and 37 °C. Data represent means ± SD, n=3. **D,** OR gene expression in plants at 22 °C or transfer to 37 °C at the time points indicated. Data are means ± SD, n=5. **E,** *OR* promoter activity before and after transfer to 37 °C in the stable transgenic lines with *OR* promoter-driven luciferase reporter. **F,** Immunoblot of OR Protein in two-week-old WT plants growing at 22 °C or transfer to 37 °C for 3 hrs. The loading amount of each sample was indicated by Ponceau S staining of the membrane and immunoblot of Actin protein. **G,** Representative phenotype of control (M82) and *OR* over-expressing (OE#8 and OE#23) tomato plants after heat stress treatment at 45 °C for 24 hrs. Anthocyanin was extracted from leaves to indicate plant stress. **H,** Analysis of chlorophyll (Chl) and carotenoid (Car) pigments with normal growth condition at 25 °C and under heat stress treatment at 45 °C for 24 hrs. Data represent means ± SD, n=3. Letters above histograms indicate significant differences as determined by Tukey’s multiple comparisons test.

Intriguingly, *OR* was found to be a heat-inducible gene. Its expression was considerably up-regulated when plants were heat-treated at 37 °C (Figure 6D). Similarly, when the *OR* promoter-driven luciferase construct was transformed into Arabidopsis, the bioluminescence signal in the transgenic lines was significantly induced when plants transferred from 22 °C to 37 °C (Figure 6E). Consequently, the OR protein level was enhanced under heat stress (Figure 6F), which promoted CHLI and PSY protein stability and safeguarded photosynthetic pigment synthesis.

Since OR is a highly conserved protein in green plant (Supplemental Figure S10), we further tested whether overexpression of *OR* has a similar effect in other plant. Two tomato *OR* overexpression lines generated previously (Yazdani et al., 2019) together with the M82 control line were treated at 45 °C for 24 hrs at the four-leaf growth stage. After the heat stress treatment, M82 was more stressed compared to the *OE* lines as visualized by more anthocyanin accumulation (Figure 6G). Furthermore, consistent with the observation in Arabidopsis, pigment analysis revealed that the tomato *OE* lines showed higher content of photosynthetic pigments than M82 under the heat stress treatment, whereas they had comparable levels with M82 control at 25 °C (Figure 6H). Together, we conclude that OR increases the resilience of photosynthesis under heat stress conditions and therefore confers plant thermotolerance.

## Discussion

The photosynthetic pigments chlorophyll and carotenoid are essential for photosynthesis, chloroplast development, and plant survival. Their biosynthesis must be coordinately regulated to efficiently and precisely adjust photosynthesis and plant fitness in response to environmental and developmental cues. However, little is known about the molecular mechanisms underlying the highly coordinated processes. Here, we discovered that the evolutionarily conserved OR proteins are the key regulators to coordinate chlorophyll and carotenoid biosynthesis. We found that OR family proteins physically interact with CHLI and regulate its protein stability for chlorophyll biosynthesis, in addition to regulating PSY for carotenoid biosynthesis (Zhou et al., 2015). Thus, OR proteins serve as common posttranslational regulators to simultaneously control the first committed enzymes of the two pathways. Moreover, OR regulates MgCh and PSY activity to safeguard photosynthetic pigment level and enhances plant thermotolerance under heat stress, providing a potential genetic target and a novel agronomic tool to generate climate-resilient crops.

Various posttranslational mechanisms are known to regulate chlorophyll or carotenoid biosynthesis (Ruiz-Sola and Rodríguez-Concepción, 2012; Brzezowski et al., 2015; Sun and Li, 2020; Wang and Grimm, 2021). The regulatory machinery via protein-protein interaction is fundamentally important to maintain and fine-tune photosynthetic pigment synthesis in chloroplast. For instance, GluTR stability is greatly enhanced by interacting with the molecular chaperone cpSRP43 (Wang et al., 2018; Ji et al., 2021) and GluTR-binding protein (Apitz et al., 2016). BALANCE of CHLOROPHYLL METABOLISM 1 (BCM1) or Chlorophyll Biosynthetic Defect1 (CBD1) was recently revealed to interact with GUN4 and synergistically function with CHLH/GUN5 to regulate MgCh activity for chlorophyll biosynthesis (Wang et al., 2020; Zhang et al., 2020). ELIP2 negatively alters the accumulation of MgCh subunits CHLI and CHLH to mediate MgCh activity for chlorophyll biosynthesis under light-stress conditions (Tzvetkova-Chevolleau et al., 2007). LIL3, a light-harvesting complex-like protein, physically interacts with POR and geranyl-geranyl reductase to coordinate chlorin and terpenoid biosynthesis by providing substrates for the subsequent step of chlorophyll *a* synthesis (Hey et al., 2017).

Compared to the regulation of chlorophyll biosynthesis, less is known about the posttranslational regulation of carotenoid biosynthesis. Deoxyxylulose 5-phosphate synthase (DXS) supplies precursors for carotenoid biosynthesis. Its activity and proteostasis are fine-regulated by interactions among several chaperone proteins (Pulido et al., 2013; Pulido et al., 2016). Previously, we showed that OR physically interacts with PSY to mediate its stability and activity for carotenoid biosynthesis in Arabidopsis and melon (Zhou et al., 2015; Chayut et al., 2017). Excitingly, we found here that OR and ORL also directly interact with CHLI, promote its folding, and stabilize the enzyme for chlorophyll synthesis. Moreover, the levels of OR family proteins positively correlated with the stability of both CHLI and PSY. Thus, through the direct interaction and maintenance of the stability of CHLI and PSY, OR family proteins coordinately regulate both pathways for photosynthetic pigment biosynthesis in plant.

The MgProtoIX level was significantly reduced in *or orl* due to impaired MgCh activity. Notably, the other intermediates including the precursor ALA and heme branch were also decreased. It has been commonly noticed that impaired MgCh activity results in low ALA biosynthesis, most likely due to an unknown feedback control (Papenbrock et al., 2000b; Papenbrock et al., 2000a). The reduction of ALA biosynthesis also reduces the content of heme as shown inter alia (Nagai et al., 2007). The reduced MgProtoIX level in *or orl* resulted in parallel or simultaneous lower steady state levels of other intermediates and consequently in decreased chlorophyll content.

OR belongs to a group of DnaJE chaperones, lacking the J-domain and only containing a C-terminal DnaJ type zinc-finger domain (Lu et al., 2006; Tzuri et al., 2015). Several members of DnaJE proteins are involved in the homeostasis or biogenesis of photosynthetic proteins or complexes. Bundle Sheath Defective 2 (BSD2) is required for ribulose-1,5-bisphosphate carboxylase/oxygenase (RuBisCo) assembly and activation (Brutnell et al., 1999). Other DnaJE chaperones are required for the PSI and PSII accumulation (Lu et al., 2011; Fristedt et al., 2014) and thylakoid membrane biogenesis (Tanz et al., 2012; Hartings et al., 2017). OR proteins stabilize CHLI and PSY in two different pathways of photosynthetic pigment synthesis. They also participate in plastid development through interactions with different partners (Sun et al., 2019; Sun et al., 2020; Yuan et al., 2021). Recently, OR family proteins have also been found to interact with photosystem components including PsbP in sweetpotato (Kang et al., 2017) and LHCB1 in melon (Chayut et al., 2021), implying their additional role in protecting photosystems other than photosynthetic pigment biosynthesis. Although the evolutionary trajectory of DnaJE proteins still needs to be established, the emergence of these DnaJE proteins in green lineage implies the acquired additional abilities to protect photosynthesis and chloroplast from hazardous environments especially during the geologic history of the earth.

Increasing the robustness of photosynthesis is a promising strategy to secure our food system since global warming has already resulted in declined crop production world-wide (Lobell et al., 2011). Reduced photosynthetic pigment accumulation or leaf chlorosis with extreme or prolonged heat stress is commonly observed in plant, reducing its overall photosynthetic capacity. It has been demonstrated that CHLI protein stability and photosynthetic pigment accumulation are significantly reduced at elevated temperatures (Rocco et al., 2013; Echevarría-Zomeño et al., 2016). Notably, the *or orl* mutant developed more severe leaf chlorosis after heat stress treatment. In contrast, plants overexpressing *OR* were less affected at elevated temperatures. This results from the role of OR family proteins in protecting plant from impairment of chlorophyll and carotenoid biosynthesis via enhancing CHLI and PSY enzyme stability. As such, OR conferred plant thermotolerance in both Arabidopsis and tomato plants, consistent with previous reports in other crops (Park et al., 2016; Kang et al., 2017). Similarly, other chaperones such as CDJ2 and RAF1 devoted to increase the efficiency and robustness of Rubisco activity also improve plant thermotolerance (Salesse-Smith et al., 2018; Ji et al., 2021).

The model of OR family proteins in coordinately regulating chlorophyll and carotenoid photosynthetic pigment synthesis and plant fitness is illustrated in Figure 7. OR and ORL are highly conserved proteins and present in all land plant. Thus, the findings uncover a common mechanism in plant to coordinate photosynthetic pigment synthesis and reveal a potential strategy to improve the resilience of heat stress in crops to address the challenge of global climate change and secure our food system.

**Figure 7.**
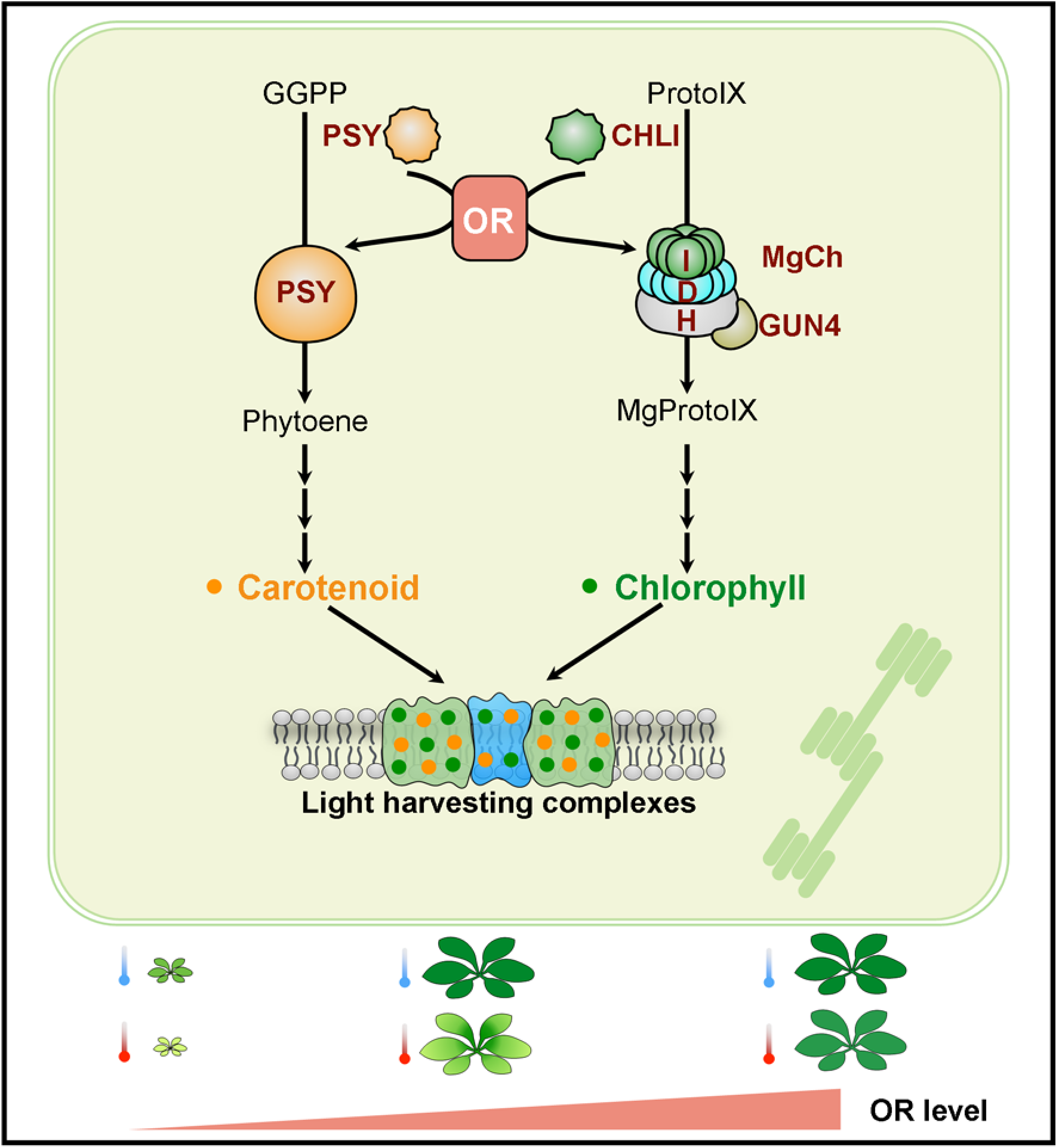
Model of OR family proteins in orchestrating both chlorophyll and carotenoid biosynthesis in chloroplast. OR family proteins directly interact with PSY and CHLI, the key enzymes for the first committed step of carotenoid and chlorophyll biosynthesis, respectively. Through the direct interactions, OR proteins mediate the stability of PSY and CHLI to coordinate photosynthetic pigment synthesis. Deficiency in OR severely impairs both chlorophyll and carotenoid biosynthesis, leading to the defect of light-harvesting complex assembly and photosynthetic thylakoid membrane stacking. OR proteins safeguard the enzyme activities of PSY and MgCh for pigment synthesis, providing a potential strategy to increase the resilience of plants to elevated temperature.

## Materials and methods

### Plant material and growth conditions

*Arabidopsis thaliana* (Arabidopsis) mutant lines including *chli* (CS44), *gun5* (CS6499), *crd1* (CS871461), and *cao* (CS41) were ordered from the Arabidopsis Biological Resource Center (ABRC). The *or orl* double mutant was generated by crossing *or* (GK-850E02-025840, GABI) and *orl* (SAIL_757_G09, ABRC) (Zhou et al., 2015). The *OR* overexpression lines in Arabidopsis and tomato (*Solanum lycopersicum*) were generated as described previously (Yuan et al., 2015; Yazdani et al., 2019). The Arabidopsis and tomato lines, including Arabidopsis WT (Columbia ecotype, Col-0) and tomato M82 wild type controls, were grown in the growth chamber with 14 h light/10 h dark cycle unless specified. The temperature was controlled at 22 °C or 25 °C for standard growth and at 37 or 45 °C for heat stress treatment for Arabidopsis or tomato.

### HPLC analysis of chlorophyll and carotenoid metabolites and heme

Analysis of chlorophyll, its precursors, and heme was performed essentially following the methods as described (Wang et al., 2020). Briefly, leaf samples were harvested from 18-day-old plants grown under short-day conditions (10 h light/14 h dark, 100 μmol photon m^−2^ s^−1^) and extracted in ice-cold acetone:0.2 M NH4OH (9:1, v/v). Following centrifugation at 14,000 *g* for 20 min at 4 °C, chlorophyll and its precursors in the supernatant were analyzed directly. Non-covalently bound heme in the pellet was extracted in acetone:hydrochloric acid: dimethylsulfoxide (10:0.5:2, v/v/v) buffer before analysis. Chlorophyll, chlorophyll precursors, and heme were analyzed by HPLC using an Agilent 1100 HPLC system equipped with a diode array and fluorescence detectors.

Analysis of carotenoid was carried out as described (Norris et al., 1995). The powdered leaf samples (50 mg) were ground in 400 μl of 80% acetone. Ethyl acetate and water (200 μl each) were added and mixed well before centrifugation at 12,000 g for 10 min. The upper phase was dried in a SpeedVac and resuspended in 100 μl ethyl acetate before analysis using Acquity UPC^2^ HSS C18 SB 1.8 mm column (3.0 × 100 mm) in the Waters UPC^2^ system (Yazdani et al., 2019). Individual carotenoid was identified using authentic standard. Quantification was carried by comparing peak area to a 5-point calibration curve of β-carotene standard and reported as β-carotene equivalent.

### Visualization of chlorophyll intermediate metabolites

The *in planta* accumulation of chlorophyll biosynthetic intermediates was visualized following the method as reported (Ankele et al., 2007) with minor modifications. Seedlings were grown on half strength MS agar medium containing 2% sucrose. To enhance the intermediate accumulation, the 6-day old seedlings were fed with ALA solution containing 10 mM ALA, 5 mM MgCl2, and 10 mM KPO_4_ (pH 7.0) in the dark for 13 h. Confocal images were acquired from 550 to 710 nm emission with the 405 nm laser excitation using an SP5 confocal laser scanning system (Leica). The emission spectra were subsequently calculated by averaging signals from 10 positions in leaves and normalized the maximal spectral values to 1. The specific emission ranges of Mg-porphyrins including MgProtoIX and MgPMME (585 to 615 nm), ProtoIX (627 to 640 nm), and chlorophyll (660 to 710 nm) were labeled as reported (Ankele et al., 2007).

### Yeast two-hybrid (Y2H) assay

Since both OR (At5g61670) and ORL (At5g06130) contain transmembrane domains, the split ubiquitin system was used for Y2H assay as described previously (Zhou et al., 2015; Welsch et al., 2018). The coding DNA sequences without transit peptides of *CHLI1, CHLI2, CHLD, GUN4, CHLH, CHLM, CRD1, PORB*, and *CAO* were cloned to make Nub plasmids. The OR-Cub plasmid was generated previously (Zhou et al., 2015). The cDNA sequence of *ORL* without transit peptide was cloned into pMetYCgate to make the Cub plasmid. Constructs were then transformed into yeast strain THY.AP4 (for Nub constructs) or THY.AP5 (for Cub constructs). Interactions were examined by mating Nub- and Cub-harboring strains and growing the mated diploid yeast colonies on a drop-out medium without leucine and tryptophan (-LW) for growth control and lacking leucine, tryptophan, adenine, and histidine (–LWAH) with 0.5 mM 3-amino-1,2,4-triazole (3-AT) for selection at 30°C for 3 days.

### Bimolecular fluorescence complementation (BiFC) assay

The BiFC assay was performed as described previously (Sun et al., 2019; Sun et al., 2020). The full-length cDNA fragments of *OR* and *ORL* and chlorophyll genes (*CHLI, CHLD, GUN4, CHLH, CHLM, CRD1*, and *PORB*) were cloned to Gateway compatible BiFC vectors pSAT4A-nEYFP-N and pSAT1A-cEYFP-N, separately. The full-length cDNA of *OR* was also cloned to pA7-YFP to get an YFP fusion. Protoplasts were isolated from Arabidopsis leaves and transfected with the N-terminal part of YFP (nY) and C-terminal part of YFP (cY) plasmid pairs by PEG mediated transfection (Yoo et al., 2007). After incubation for 16 h, the protoplasts were observed under a SP5 Laser Scanning Confocal Microscope (Leica) with the laser excitation wavelength at 488 nm. The YFP fluorescent signal was detected through an emission filter between 520 nm and 560 nm. The chlorophyll fluorescent signal in chloroplast was detected between 660 to 710 nm.

### Blue-Native PAGE, SDS-PAGE, and immunoblotting

Blue native (BN)-PAGE was carried out as described (Zhou et al., 2017) with minor modifications. Leaves of WT and *or orl* mutant were homogenized in ice-cold isolation buffer containing 330 mM sorbitol, 50 mM HEPES-KOH (pH 7.5), 2 mM EDTA, 1 mM MgCl_2_, 5 mM ascorbate, 0.05% bovine serum albumin, and 1% protease inhibitor cocktail (Sigma-Aldrich). The suspension was filtered through 100 μm and 40 μm nylon and centrifuged at 1,000 g at 4°C for 5 min. The pellet was resuspended in the isolation buffer and loaded on a two-step Percoll gradient with 80% (w/v) bottom and 40% (w/v) top in isolation buffer. The samples were centrifuged in a swing-out rotor at 4,000 *g* for 10 min with a brake off setting. The band appeared between two phases was piped out and washed twice with wash buffer (50 mM HEPES, 3 mM MgSO_4_, and 0.3 M sorbitol). Solubilization of chloroplast was performed by addition of 1% (w/v) Triton X-100 and incubation on ice for 20 min.

BN-PAGE was performed using Novex NativePAGE (Life Technologies) 4-16% acrylamide gradient gels according to the manufacturer’s protocol. For each lane, a protein sample containing 40 μg of chlorophyll was loaded. The major colored thylakoid membrane complexes on BN-PAGE gel were estimated as described before (Wang and Grimm, 2016). For the second dimension SDS-PAGE and immunoblot, duplicate gel strips from BN-PAGE were excised and incubated with 1% SDS and 1% β-mercaptoethanol at 37 °C for 30 min. The treated gel strips were then washed with water and placed on top of 15% SDS-PAGE gels. After electrophoresis, proteins were blotted onto nitrocellulose membrane (Millipore) and probed with corresponding antibodies.

For immunoblot analysis of leaf total protein samples, proteins were extracted by homogenization in protein extraction buffer containing 100 mM Tris-HCl (pH 6.8), 4% sodium dodecyl sulfate (SDS), 1 mM dithiothreitol (DTT), 1 mM phenylmethylsulfonyl fluoride (PMSF) and 1 x Protease Inhibitor Cocktail (NEB) as described before (Sun et al., 2019). Proteins were separated by 15% SDS-PAGE gel and blotted onto a nitrocellulose membrane (Millipore). The membrane was first stained with Ponceau S to show protein loading before blocking and incubation with first antibodies: CHLI antiserum was raised by Genscript (Nanjing, China) and validated in *chli* mutant and WT (Supplemental Figure S12); PSY antibody was generated by Abgent (San Diego, CA); CRD1 antibody was purchased from PhytoAB (Cat# PHY1312S); D1 (Cat#AS05084), PORB (Cat#AS05067), LHCA1-4, and LHCB1-6 (Cat#AS01011) antibodies were purchased from Agrisera. A secondary goat-anti-rabbit HRP-conjugated antibody (BioRad cat#1706515) was used with the dilution of 1:10000 and a WesternBright ECL kit was used to detect the chemiluminescent signals (LPS Cat# K-12045-D20).

### Proteomics analysis

Total proteins from leaves of WT, *OE* line, and *or orl* with three biological replicates were extracted using the phenol extraction method as described (Yang et al., 2007). The subsequent proteolytic digestion and labeling using TMT 10-plex reagents were carried out according to the manufacturer’s recommended protocol (Thermo Scientific, MA, USA). The TMT 10-plex tagged tryptic peptides were fractionated using high pH reversed phase (hpRP) chromatography as reported previously (Yang et al., 2011). The subsequent Nano LC-MS/MS analysis was performed on an Orbitrap Elite mass spectrometer (Thermo-Fisher Scientific, San Jose, CA) coupled with the UltiMate3000 RSLCnano (Dionex, Sunnyvale, CA) with high energy collision dissociation (HCD). The MS/MS data was acquired by Xcalibur 2.2 software (Thermo-Fisher Scientific) and searched against the Arabidopsis database (https://www.arabidopsis.org/download/index-auto.jsp?dir=/download_files/Proteins). TMT quantification was performed using Sequest HT software within the Proteome Discoverer 2.2 (PD2.2). All reporter ions designated as “quantifiable spectra” were used for peptide/protein quantitation.

### Transmission electron microscopy (TEM)

For TEM observation of chloroplast ultrastructure, 3-week-old plant leaves were fixed, embedded, and sectioned as described previously (Sun et al., 2019). Briefly, leaves were first harvested and kept in 4.0% (w/v) formaldehyde and 2.5% (w/v) glutaraldehyde in 0.1 M phosphate buffer (pH 7.2). They were then fixed with 1% (w/v) osmium tetroxide in 0.1 M phosphate buffer (pH 7.2) containing 1.5% (w/v) potassium ferricyanide. Samples were subsequently embedded in Spurr’s low-viscosity resin. After resin polymerization, the embedded samples were cut into ultrathin sections. They were stained in aqueous uranyl acetate and Reynolds lead citrate, and picked on copper grids before observation. Image captures were carried out on a Hitachi-7650 transmission electron microscope.

### Fv/Fm measurement

For Fv/Fm measurement of WT, *or orl*, and *OE* lines, the initial determination of F0 and Fm was done by the application of a saturation pulse (2,800 μmol photons m^−2^ s^−1^) after 20 min of dark adaptation using FluorCam as described (Murchie and Lawson, 2013). Fv/Fm was calculated using the following equations: Fv/Fm = (Fm – F0)/Fm.

### Expression and purification of recombinant proteins

The coding sequence of *CHLI* without transit peptide (1-46 amino acid residues) was cloned into pET32a (Novagen) and pGEX-4T-1 (GE healthcare) vectors for prokaryotic expression in *E. coli* BL21(DE3)pLysS (Novagen). The coding sequences for the N-terminal (63-139 aa) and C-terminal (221-307 aa) domain of *OR* were cloned to the pGEX-4T-1 vector (GE healthcare) for prokaryotic expression. His-tagged and GST-tagged recombinant proteins were purified using Promega MagneHis™ Ni-Particles (Promega Cat#V854A) and MagneGST™ Glutathione Particles (Promega Cat# V861A) following the manufacturer’s manual. Expression and purification of His-tagged MgCh subunits from *Oryza sativa* (OsCHLH, OsCHLD, and OsCHLI) and GUN4 were reported previously (Ji et al., 2021). All the primers used for cloning are shown in Supplemental Table S1.

### Gene expression analysis

Gene expression analysis was carried out as described previously (Yuan et al., 2021). Total RNA from leaves of WT, *or orl*, and *OE* lines was extracted using TRIzol Reagent (Invitrogen) and reverse transcribed into cDNA. RT-qPCR was performed on a CFX384 Touch Real-time PCR Detection System (Bio-Rad) using SYBR Green Master Mix (Bio-Rad) and gene-specific primers (Supplemental Table S1) with *Actin 8* and *UBQ* as internal controls. The analysis was carried out with three biological replicates and three technical trials.

### Luciferase reporter assay

The ORpro:LUC construct was obtained from our previous study (Sun et al., 2019). Basically, the Luc+ gene from pSP-Luc+NF (Promega) was cloned into pCAMBIA1390 (CAMBIA) to generate pCAMBIA1390-Luc+. A 2000-bp upstream flanking region of OR was then inserted to drive the expression of luciferase. The construct was electroporated into GV3101 and transformed into Arabidopsis plants by floral dipping. Five independent T3 homozygote lines were used for heat treatment on half MS agar plates with 100 μg ml^−1^ luciferin. The bioluminescence of plants was detected and quantified by the BioRad ChemiDoc MP imaging system before and after heat stress treatment at 37 °C for 3 hrs.

### *In vitro* refolding assay

The refolding assay was carried out following the reported method (Wang et al., 2018) with minor modifications. Bacteria harboring His-tagged or GST-tagged CHLI protein were treated at 42 °C for 1 hour to increase aggregated CHLI protein fraction. The bacterial cells were lysed and pelleted to get the insoluble fraction of CHLI. The pellet was resuspended and incubated with GST, GST-OR-N, and GST-OR-C proteins separately. After incubation at room temperature for 1 hr, these samples were centrifuged at 15,000 *g* for 10 min to separate soluble and pellet fractions. Both fractions were adjusted to the same volume with 1X SDS Loading buffer and equal volume (10 μL) of each sample was loaded onto SDS-PAGE gel for further immunoblot against CHLI. After immunoblot, band intensities were quantified three times using ImageJ and the percentage of soluble and pellet fractions of CHLI was calculated.

### *In vitro* MgCh activity assay

The *in vitro* MgCh activity assay was performed as described previously (Ji et al., 2021) with minor modifications. The reaction mixture of 150 μL contained 2.5 μM CHLH, 1 μM CHLD, 1 μM CHLI, and 2.5 μM GUN4. CHLH, GUN4, and 5 μM ProtoIX were pre-incubated with 2.5 μM GST, GST-OR-N or GST-OR-C in 100 μL of MgCh assay buffer containing 50 mM Tricine-KOH, pH 8.0, 15 mM MgCl2, 0.2 mM DTT, and 1 mM ATP in darkness for 30 min on ice. Separately, CHLD and CHLI were pre-incubated in 50 μL of the MgCh assay buffer. The two mixtures were then gently combined and incubated at 30 °C for 5 min or at 42 °C for 5 min. The production of MgProtoIX was continuously monitored by spectrofluorometry using a 96-well plate reader mounted on a Hitachi F7000 fluorescence photometer. A standard curve was used to calculate MgProtoIX amounts. The protein levels at the end of the reaction were checked with 12% SDS-PAGE.

## Supplemental Data

**Supplemental Table S1**. Primers used in this study

**Supplemental Table S2**. Proteomic profiling of differentially expressed photosystem proteins in *or orl* compared to wild type control

**Supplemental Figure S1**. Interaction between OR family proteins and chlorophyll biosynthetic enzymes by Y2H

**Supplemental Figure S2**. Interaction between OR family proteins and chlorophyll biosynthetic enzymes by BiFC

**Supplemental Figure S3**. Sub-organelle localization of OR, PSY, and CHLI proteins

**Supplemental Figure S4**. Staining of the BN-PAGE and SDS-PAGE gel

**Supplemental Figure S5**. Steady state levels of light harvest complex proteins

**Supplemental Figure S6**. Refolding assay of GST-CHLI and His-CHLI proteins

**Supplemental Figure S7**. SDS-PAGE analysis of recombinant proteins used in MgCh activity assay

**Supplemental Figure S8**. *In vitro* MgCh activity assay at 22 °C by the measurement of MgProtoIX fluorescence intensity

**Supplemental Figure S9**. OR protein level in WT, *or orl* and *OE* lines by immunoblot analysis

**Supplemental Figure S10**. Phylogenetic tree of OR family proteins in green linage

**Supplemental Figure S11**. Validation of CHLI antibody in wild type and *chli* mutant

## Author contributions

TS and LL designed the research. TS performed the majority of the experiments. PW carried out chlorophyll intermediate metabolite analysis and *in vitro* MgCh activity assay. SL provided electron microscopy analysis. HY generated some genetic materials. YY and TF did proteomics and assisted carotenoid analysis. TS, PW, BG, and LL analyzed the data. SL, TWT, JL, MM, and BG contributed research agents, assisted data interpretation, and revised the manuscript. TS and LL wrote the article with contributions from all coauthors.

